# UK BioCoin: Swift Trait-Specific Summary Statistics Regression for UK Biobank

**DOI:** 10.1101/2024.04.12.589273

**Authors:** Jing-Cheng He, Guo-An Qi, Jiacheng Ying, Yu Qian, Lide Han, Yingying Mao, Hou-Feng Zheng, Hangjin Jiang, Guo-Bo Chen

## Abstract

Summary statistics derived from large-scale biobanks facilitate the sharing of genetic discoveries while minimizing the risk of compromising individual-level data privacy. However, these summary statistics, such as those from the UK Biobank (UKB) provided by Neale’s lab, are often adjusted by a fixed set of covariates to all traits (12 covariates including 10 PCs, sex and age), preventing the exploration of trait-specific summary statistics. In this study, we present a novel computational device UK BioCoin (**UKC**), which is designed to provide an efficient framework for trait-specific adjustment for covariates. Without requiring access to individual-level data from UKB, UKC leverages summary statistics regression technique and resources from UKB (289 GB of 199 phenotypes and 10 million SNPs), to enable the generation of GWAS summary statistics adjusted by user-specified covariates. Through a comprehensive analysis of height under trait-specific adjustments, we demonstrate that the GWAS summary statistics generated by UKC closely mirror those generated from individual-level UKB GWAS (*ρ ≥* 0.99 for effect sizes and *ρ ≥* 0.99 for *p*-values). Furthermore, we demonstrate the results for GWAS, SNP-heritability estimation, polygenic score, and Mendelian randomization, after various trait-specific covariate adjustments as allowed by UKC, indicating UKC a platform that harnesses in-depth exploration for researchers lacking access to UKB. The whole framework of UKC is portable for other biobank, as demonstrated in Westlake Biobank, which can equivalently be converted to a ‘UKC-like” platform and promote data sharing. UKC has its computational engine fully optimized, and the computational efficiency of UKC is about 70 times faster than that of UKB. We package UKC as a Docker image of 20 GB (https://github.com/Ttttt47/UKBioCoin), which can be easily deployed on an average computer (e.g. laptop).

**One sentence summary:** We develop UK BioCoin (UKC), which allows fine-tuning of covariates for each UK Biobank trait but does not relay on UK Biobank individual-level data. It will change the current landscape of GWAS and reshape its downstream analyses.

## 1 Introduction

Summary statistics, including estimated allelic effect sizes, standard errors of the estimates and other per-SNP features, are increasingly generated from genome-wide association studies (GWAS) across thousands of human traits [1, 2]. Compared to individual-level data, summary statistics raise fewer privacy concerns, making them a useful intermediary for data-sharing. The availability of publicly accessible summary statistics databases is expanding, in response to the growing demand for reproducibility and follow-up analysis of GWAS results [3]. The utility of summary statistics, including meta-analysis, gene-based association analysis, polygenic prediction, and more, provides insights of genetic architecture of complex human traits, particularly through large-scale collaborations among biobanks [4, 2, 5].

However, the current data-sharing mode based on summary statistics has several limitations. While it is common practice to adjust for covariates such as sex and age in GWAS, there is no universally applicable set of covariates for all traits, and inappropriately chosen covariates may reduce the power of findings and even introduce bias when they act as confounders [6]. For example, UK Biobank (UKB) is one of the most cited data sources for GWAS [7, 8], and the available UKB GWAS summary statistics are trained under a predefined model, such as released by Neale’s Lab (by adjusting 10 principal components, sex, and age; https://nealelab.github.io). As demonstrated in our study of UKB data, the inclusion or exclusion of certain covariates can lead to significantly different summary statistics, thereby influencing downstream analyses. An ideal summary statistics analysis framework may permit efficient in-depth explorations of different covariates setups for each trait. However, refinement of covariates is cumbersome and time-consuming for large-scale collaboration, which usually involves several rounds of rerunning GWAS at up to dozens of different biobanks [2, 5], highlighting the urgent need for a more efficient engine to generate GWAS summary statistics.

In this study, we propose a novel framework for summary statistics sharing and presents a working instance called UK BioCoin (UKC, herein) corresponding to UKB, targeting both trait-specific and efficient generation of summary statistics. The UKC framework promises highly efficient trait-specific covariates exploration while maintaining the data-sharing virtue of summary statistics, thereby promoting collaborations, especially in the context of large-scale biobank studies.

As demonstrated, summary statistics generated from UKC and the individual-level UKB is nearly identical or practically consistent across a serial of models. Furthermore, the UKC computational kernel reduces computational time complexity by nearly two orders compared to the UKB GWAS conducted in PLINK2 (PLINK herein) [9], and this efficiency significantly facilitates the exploration of competitive GWAS models and increases the robustness of a study even for researchers who do not directly access UKB resources. The whole framework of UKC is comprehensively illustrated using UKB and can be readily applied to other biobanks, such as demonstrated in the Westlake Biobank [10].

## 2 Results

### 2.1 Sketch for UK BioCoin

In this study, we allow UKC to train a trait-specific GWAS model under the choice of different covariates, while anyone using UKC does not require to access UKB individual-level data. As a proof-of-principle study, we focus on the analysis of 292,216 unrelated individuals of white British and Irish descent in the UK Biobank (UKB Field ID 22021 and 21000). 10,531,641 quality-controlled single nucleotide polymorphisms (10M SNPs herein) are included (**Fig. 1 A**). The effective number of SNPs is about *m*_*e*_ = 161, 688, or equivalently, the genomic LD is about 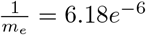. As expected, chromosomal LD is proportional to inversion of chromosome length (**Fig. 1 B**). *F*_*st*_ *≈* 0.00014 indicates little population structure among UKB samples [11]. We examine 129 conventional UKB phenotypes, comprising 60 continuous traits and 69 categorical traits. Each phenotype is scaled to have a mean of zero and a variance of one. **Fig. 1 C** illustrates the pairwise correlation between the 129 phenotypes, of which the overall missing rate is 4.1%. These 129 traits can be divided into 8 categories, such as baseline characteristics and social demographics according to the UKB catalogue, and more detailed information on these traits can be found in **Supplementary Data I**. We surrogate population structure with the top 30 principal components directly estimated from 1 million sampled SNPs from the 10M SNPs (UKC-PCs, default PCs for analysis otherwise specified); for comparison and compatibility, we also include the top 40 PCs as originally provided by UKB (UKB Field ID 22009; UKB-PCs).

**Figure 1.**
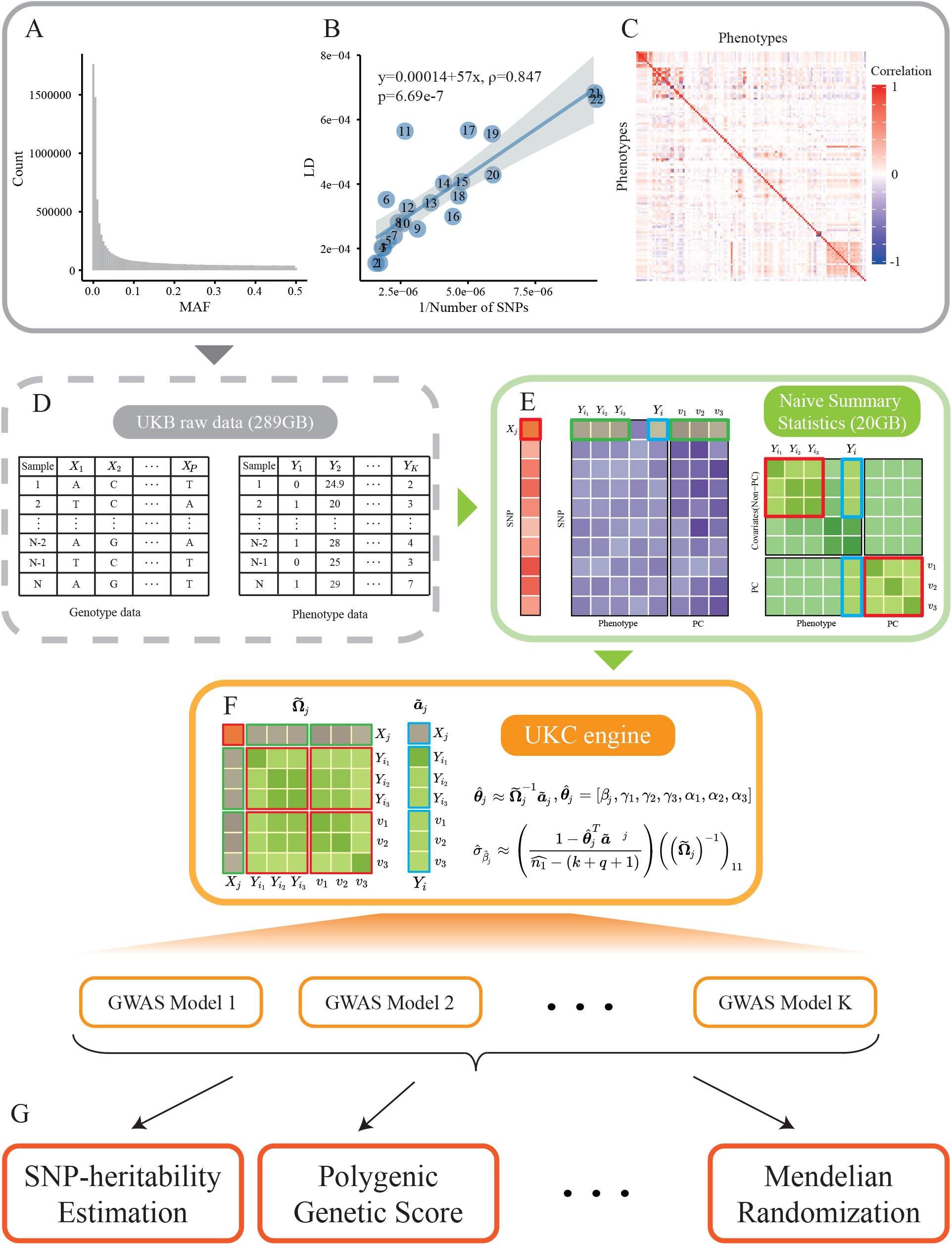
Outline of UK BioCoin and its interface to other genetics applications. **A**) The distribution of minor allele frequency of the QCed 10,531,641 SNPs included in UK BioCoin (UKC), and their MAFs are greater than 0.001. **B**) Chromosome-wise linkage disequilibrium of 22 autosomes. The fitted regression line, *y* = 0.00014 + 57*x*, indicates the linear correlation between chromosomal LD and the inversion of chromosomal length. *ρ* = 0.847 quantifies the correlation between *x* and *y*; the intercept of 0.00014 represents genomic *F*_*st*_. **C**) The correlation heatmap of 129 phenotypes used in UKC. **D-F**) UKC naive summary statistics (**E**) are derived from UKB raw data (**D**). The UKC engine (**F**) utilizes the NSS to perform regression approximately 70 times more efficient than PLINK while requiring significantly reduced memory. **G**) UKC results enable downstream genetic applications.

The UKC framework, described in **Fig. 1 D-G**, comprises two main components. **I**) The naive summary statistics (NSS) derived from UKB individual-level data. NSS is essentially a set of primary GWAS summary statistics and is consistent with the data sharing policy for UKB. **II**) A highly efficient summary statistics regression engine [12, 13]. For a GWAS model, the regression engine retrieves the required statistics from the NSS to generate trait-specific summary statistics. We evaluate the quality of the UKC results by comparing them with those of individual-level UKB data via PLINK. Compared to PLINK, UKC offers superior computational efficiency and demonstrates high consistency with PLINK, particularly when missing rates are low. Furthermore, a single quality control metric, the variation of inflation (VIF), can safeguard high-quality GWAS summary statistics (**Fig. 2**). The calculation details are provided in the **Methods** section.

**Figure 2.**
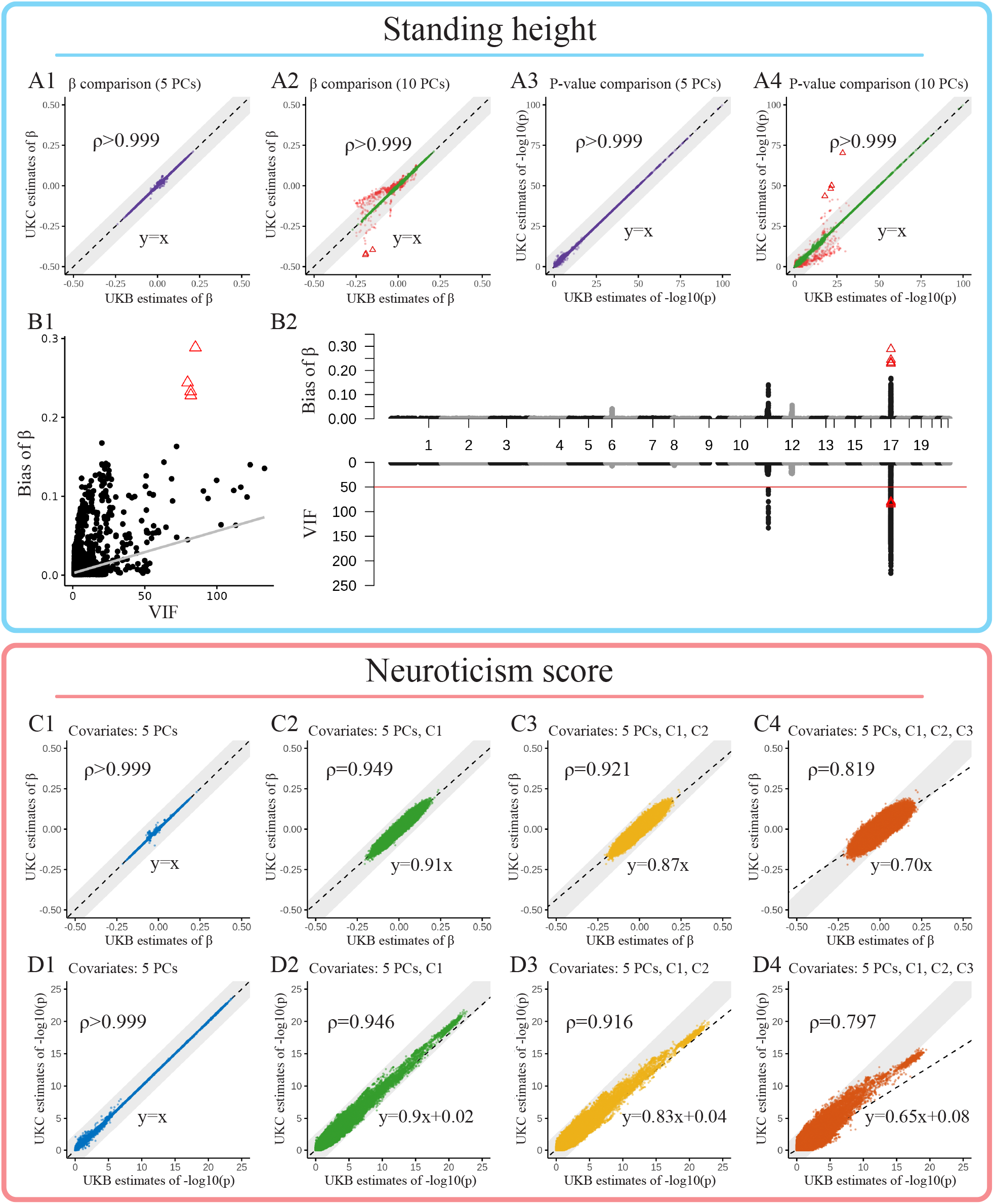
Performance of UK BioCoin comparing to UK BioBank under various adjustments. **A**) Comparison of regression coefficient (**A1-A2**) and *−*log_10_(*p*) (**A3-A4**) generated by UK BioCoin and PLINK for GWAS for **Standing height**, adjusted for 5 and 10 principal components accordingly. In the model adjusted for 10 PCs (**A2, A4**), the SNPs with 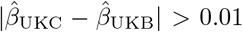 are highlighted in red, and the SNPs with 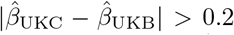 and VIF *>* 50 are labeled with triangles. **B**) Correlation (**B1**) and Miami plot (**B2**) of VIF and bias 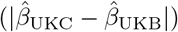. The included PCs are all UKC-PCs. **C-D**) Comparison of regression coefficient (**C**) and *−*log_10_(*p*-value) (**D**) generated by UK BioCoin and PLINK when missing rate is higher than 10%. The target phenotype is **Neuroticism score** (missing rate *≈* 18.7%), and from left to right the covariates included were: **C1 Exposure to tobacco smoke at home** (missing rate *≈* 9.3%), **C2 Snoring** (missing rate *≈* 6.8%), and **C3 Alcohol usually taken with meals** (missing rate *≈* 20.7%) is subsequently added to the model as covariates.

#### 2.1.1 Generation of Naive Summary Statistics for UK BioCoin

The generation of UKC NSS mainly involves calculating Pearson’s correlation between each SNP and each trait. This process incurs a significant computational cost, approximately *O*(*n*(*K* + *Q*)*P*) depending on the number of SNPs (*P*), phenotypes (*K*), eigenvector (*Q*), and sample size (*n*). In this study, it totals the calculation for 10*M ×* (129 + 30 + 40) Pearson’s correlation, which accounts for 129 traits, 30 UKC-PCs, and 40 UKB-PCs against each of the 10M SNPs. The main component of UKC NSS is a matrix that consequently has dimensions of 199 *×* 10*M*, effectively compressing the UKB raw data from nearly 289 GB, encompassing 129 phenotypes and approximately 10 million QCed SNPs (referred to as 10M SNPs), to less than 20 GB of NSS. The correlation between a SNP with each of the 129 traits is equivalent to estimate its effect size in a GWAS model without any adjustment, and the correlation between a SNP with UKC-PCs or UKB-PCs is known as EigenGWAS [14]. Other complementary summary statistics are generated, such as the variance of each SNP, correlation matrix between all traits, but they take much less storage and calculation than the main NSS matrix.

It takes approximately 2 days to generate UKC NSS on a cluster with 60 threads. Although it seems expensive to generate the NSS, it brings in significant efficiency in the downstream GWAS for complex traits. The details of UKC NSS generation are described in the **Methods** section.

#### 2.1.2 Computational Efficiency of UK BioCoin

The efficient performance of UKC is made possible by both algorithmic and programming advantages. The computational complexity for a linear regression is approximately *O*(*np*^2^ + *p*^3^) for a testing SNP, where *n* is the sample size and *p* is the number of covariables in a GWAS. In particular, *O*(*np*^2^) is the cost to generate the correlation matrix **Ω** of *p* variables and *O*(*p*^3^) the inversion for **Ω**. On the contrary, UKC constructs **Ω** by accessing the corresponding elements in NSS matrices, so *O*(*np*^2^) is completely dismissed. Furthermore, when UKC moves from the *i*^th^ to the *j*^th^ locus, only the first column and the first row of **Ω** are updated (purple blocks in green boxes and red block in red box in **Fig. 1 F**) and leave the submatrix **Ω**_*−*1,*−*1_ (**Ω**_*−*1,*−*1_ refers to the submatrix of **Ω** by dropping the first row and the first column, and corresponds to the green blocks in red boxes in **Fig. 1**) the same for each locus. It enables the blockwise inversion technique, and since the inversion of **Ω**_*−*1,*−*1_ is performed only once for the whole scanning of 10M SNPs, and the original *O*(*p*^3^) for **Ω**^*−*1^ is reduced to *O*(*p*^2^) for each locus. So the computational cost of a test SNP is reduced from *O*(*np*^2^ + *p*^3^) to *O*(*p*^2^).

Secondly, the UKC computational engine is implemented in C++ and uses the Eigen library for efficient and precise matrix computations [15]. UKC leverages the efficient looping capabilities of the C++ language, enabling accelerated program execution, particularly for a large-scale dataset containing millions of SNPs. UKC adopts a stream processing strategy that minimizes memory consumption by loading only a fraction of the data at any given time. Both pre-calculated NSS and advanced programming allow UKC to execute multiple tasks simultaneously and efficiently, even on a personal laptop.

We compare the efficiency of UKC and UKB in conducting the 3 GWAS models for Standing height (UKB field ID: 50) with adjustment of 0, 5, and 10 PCs, respectively. As tested, using 16 threads on a cluster, PLINK took about 3 hours to perform GWAS on 10M SNPs with 5 covariates; in contrast, UKC took 0.6 hours only using a single thread to complete the same task, a boost that improves computational efficiency about 80 times. In terms of memory usage, PLINK required approximately 5 GB of peak memory, while UKC required less than 5 MB (**Tab.1**).

**Table 1:** Comparison of computational efficiency of PLINK and UKC.

| Phenotype | Covariates | Method | Num. of threads | Running time | Memory used |
| --- | --- | --- | --- | --- | --- |
| Standing height | None | PLINK | 16 | 0.79 h | 4.89 GB |
|  |  | UKC | 1 | 0.17 h | 2.64 MB |
|  | 5 PCs | PLINK | 16 | 3.05 h | 5.11 GB |
|  |  | UKC | 1 | 0.60 h | 2.74 MB |
|  | 10 PCs | PLINK | 16 | 4.49 h | 5.29 GB |
|  |  | UKC | 1 | 0.98 h | 2.77 MB |

### 2.2 Quality Control for UK BioCoin

#### 2.2.1 Influence of Phenotype Missing Rates

UKC generates identical results to those of UKB when there is no missing data (see **Methods**). However, missing data occurs, leading to differences in **Ω** of different degree, and possibly introduces noise to UKC. We incorporated 0, 5, and 10 PCs as the covariates for Standing height (UKB field ID: 50, of low missing rate *<* 1%) for UKC, and for comparison an identical UKB model was then performed in PLINK. We compared the SNP effects (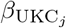 and 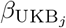, and defined bias 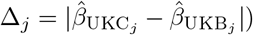 and their corresponding *p*-values between UKC and UKB, and for all three GWASs their respect Pearson’s correlation was greater than 0.999 (**Fig. 2 A1-A4**). Remarkably, in all 3 GWASs, UKC recovered *>* 99% significant SNPs (*p*-value *<*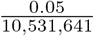) as found in UKB (**Supplementary Data II**). We further decomposed the difference for the *j*^th^ locus Δ_*j*_ = *δ*_*j*_ *·* VIF_*j*_. When the model was adjusted by 10 PCs, 1,104 inconsistency SNPs had Δ_*j*_ *>* 0.01 on chromosomes 6 (HLA cluster), 11, 12, and 17 (red points in **Fig. 2 A2, A4, B1-B2**), and all these SNPs had high VIF; in particular, severe inconsistency (Δ_*j*_ *>* 0.2) was associated with extremely high VIF (VIF_*j*_ *>* 50, red triangles in **Fig. 2 A2, A4, B1-B2**). In this example, the inclusion of too many covariates such as PCs was likely to lead to high VIF, which amplified bias. As PCs were orthogonal to each other, we could derive an analytical result, **Eq** 14 in **Methods**, which characterized how biased SNPs were and how their effects were further amplified by VIF. To minimize biases introduced by approximation in the UKC, one could use a stringent VIF threshold. Excluding the SNPs with VIF *>* 50, as default in PLINK, removed those severe inconsistent loci (Δ_*j*_ *>* 0.2). Few SNPs had high VIF and that even adopting VIF *>* 10 as cutoff only removed less than 0.1% of the 10M SNPs in the model with 10 PCs.

Furthermore, we directly examined UKC under exceptionally high missing rates. In this experiment, the phenotype was Neuroticism score (UKB field ID: 20127, missing rate of 18.7%) and was adjusted by the top five PCs and three covariates of high missing covariates: Exposure to tobacco smoke at home (UKB field ID: 1269, missing rate of 9.3%), Snoring (UKB field ID: 1210, missing rate of 6.8%), and Alcohol usually taken with meals (UKB field ID: 1618, missing rate of 20.7%). When incorporating additional covariates, the inconsistency between 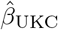 and 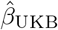 increased, suggesting that the missing pattern of phenotypes included in the model was non-random (**Fig. 2**), and the lowest correlations for *β* and log_10_(*p*) were 0.819 and 0.797 respectively. In general, although UKC produced more conservative estimates when the missing rate was high (**Fig. 2 C2-C4, D2-D4**), the significant genetic variants identified by UKC and UKB were generally consistent. The details of the results are given in **Supplementary Data II**. To benchmark the influence of missing data, we randomly sampled a phenotype and 3 covariates from the 129 traits, and its identical model was also analyzed using UKB data with PLINK. We repeated this procedure 50 times, and top 5 PCs were always included in a model. The consequent correlation for 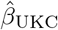 and 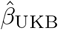 was 0.937 (s.d. 0.043) for log_10_(*p*) was 0.901 (s.d. 0.068), respectively. So the influence of missing rate on average was less severe than the Neuroticism score example.

In general, UKC reproduced the GWAS results with remarkable precision when the missing rates of phenotypes were low. In situations with high overall missing rates, estimates might exhibit conservative bias but were still closely consistent with results of individual-level data. As VIF was useful to exclude potentially misleading GWAS signals, in the analysis below, we used VIF = 50 as the default threshold to remove potentially abnormal GWAS signals. Synthesizing VIF metrics cost little because each VIF value was windfall for its testing SNP (see the **Methods** section).

### 2.3 UK BioCoin for In-depth Genetic Exploration

As illustrated in **Fig. 1**, UKC enables in-depth exploration for many genetic studies. We are going to illustrate how our UKC can be flexibly integrated into downstream genetic studies, which have GWAS summary statistics as input, and uncover the variation of these genetic studies due to trait-specific adjustment. Here, we present four typical applications of UKC: **I**) GWAS of various adjustments; **II**) SNP-heritability estimation by LD score regression (LDSC, [16]); **III**) polygenic score as generated via “–score” in PLINK [9]); **IV**) Mendelian randomization for exploring casual effects of waist circumference on rheumatoid arthritis.

#### 2.3.1 Application 1: GWAS with Flexible Covariate Adjustment

For the subject matter of the presentation, the covariates for GWAS are divided into three categories: **I**) covariates without or of little heritability but of biological significance, such as sex [17]; **II**) covariates with heritability, such as height and BMI, which are known to influence the outcome of GWAS due to genetic correlation [6, 18]; **III**) covariates for population structure, surrogated by principal components [19, 20, 21]. We demonstrate in traits Standing height and Weight (UKB field ID: 21002) how UKC provides additional information than a conventional GWAS (**Fig. 3**).

**Figure 3.**
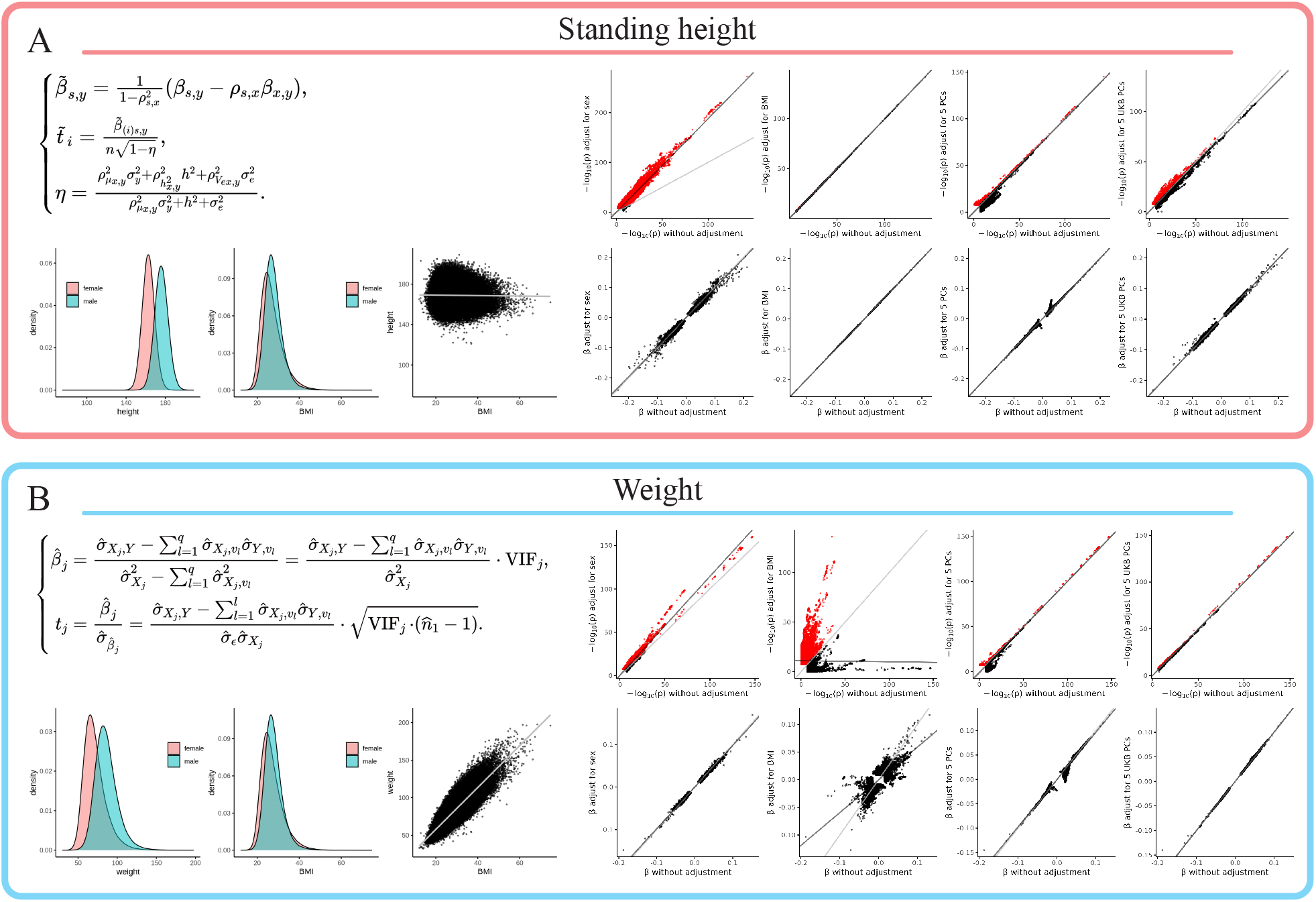
UKC conducts GWAS for Standing height and Weight under various adjustments. For each trait, the first row is for -log_10_(*p*) and the second row for *β*, in each plot x and y axes compare with and without adjustment for sex (first column), BMI (second column), 5 top UKC-PCs (third column) and 5 top UKB-PCs (forth column). Sex represents a covariate of low/no heritability, BMI a covariate of high heritabiltity, and PCs for adjustment for population structure.

Sex (UKB field ID: 31), which was obviously not associated with 10M SNPs, explained *R*^2^ *≈* 0.5 of the variation of height between men and women. With or without inclusion of Sex, the genetic effects were little changed, but with the inclusion of Sex the statistical power increased significantly and the number of associated loci increased from 47,790 to 128,730 SNPs before clumping. When Standing height was adjusted by BMI (UKB field ID: 21001), which had *h*^2^ = 0.24 itself but of little correlation with Standing height, it showed an ignorable effect of the adjustment (**Fig. 3 A**).

On the contrary, the pattern differed significantly for Weight after adjustment. After adjustment for Sex, which explained approximately *R*^2^ *≈* 0.21 for Weight, there was a slight increase in statistical power, and the estimation of *β* was negligibly influenced. However, after adjustment for BMI, which was highly correlated with Weight, statistical power was stratified for loci that influence both Weight and BMI, and in addition, the genetic effects were significantly altered. On closer examination of the results, of 47,790 SNPs significantly associated with Standing height, 47,176 remained significant with adjustment of BMI. On the contrary, of 20,912 SNPs significantly associated with Weight, only 7,450 remained significant after BMI adjustment. Although covariates with certain heritability (such as BMI) were commonly included, they were likely act as confounders in the study and would be considered to bias the effects estimates [6] (**Fig. 3 B**). It was upon the purpose of a study to justify the adjustment.

For both traits, with or without adjustment for the top 5 PCs made little difference for the estimation of *β* and their statistical power, regardless of whether the PCs were either UKC-PCs or UKB-PCs. The visible difference was observed, but only for SNPs of very small effect sizes, probably because of subtle local population structure. The detailed underlying statistical mechanism are provided in the **Methods** section. For 129 traits, we applied five adjustment schemes (no adjustment at all, 5 PCs, 10 PCs, 5 PCs with sex and 5 PCs with BMI), and their summary results are given in **Supplementary Data III**.

While using covariates without heritability may increase power, this is only true when they are not confounding factors. In some case-control studies, the ascertainment for case/control samples may create correlations between trait and covariates that are not presented in a natural population. Adjusting for these covariates could decrease power and potentially introduce bias [22]. Since UKC runs on population data rather than ascertained samples, this problem was less likely to arise. Researchers must consider covariate characteristics, such as heritability and relevance to the trait under study, to fit the purpose of their studies.

#### 2.3.2 Application 2: Estimation for SNP-heritability

One windfall of GWAS summary statistics is the estimation of SNP heritage 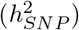 using LDSC [16]. For each of the 129 traits, UKC generated eight GWAS summary statistics, which were adjusted by i) no covariates; ii) 5 PCs; iii) 10 PCs; iv) 5 PCs and Sex; v) 5 PCs and BMI; vi) Sex only; vii) BMI only; viii) 5 PCs, Sex and Age (UKB field ID: 21022). These eight sets of GWAS summary statistics were fed into LDSC, which included HapMap3 SNP variants with MAF *>* 0.001 totaling 1.17M SNPs. For most traits, their 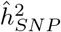 showed little variation regardless of adjustment schemes, probably because these traits had little heritability (**Fig. 4 A**), and the adjustments resulted in slight variations in the means of the heritability estimates of the 129 traits (**Fig. 4 B**). However, for traits in category “Physical measure”, especially for those with visible differences between men and women such as Standing height and Weight, inclusion or exclusion of sex as a covariate resulted in different heritability estimates. Subtle population stratification could have an impact on the estimation of heritability, as evidenced by a significant increase 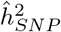 of Weight after correcting for 10 PCs. A complete summary table of the results is provided in **Supplementary Data IV**.

**Figure 4.**
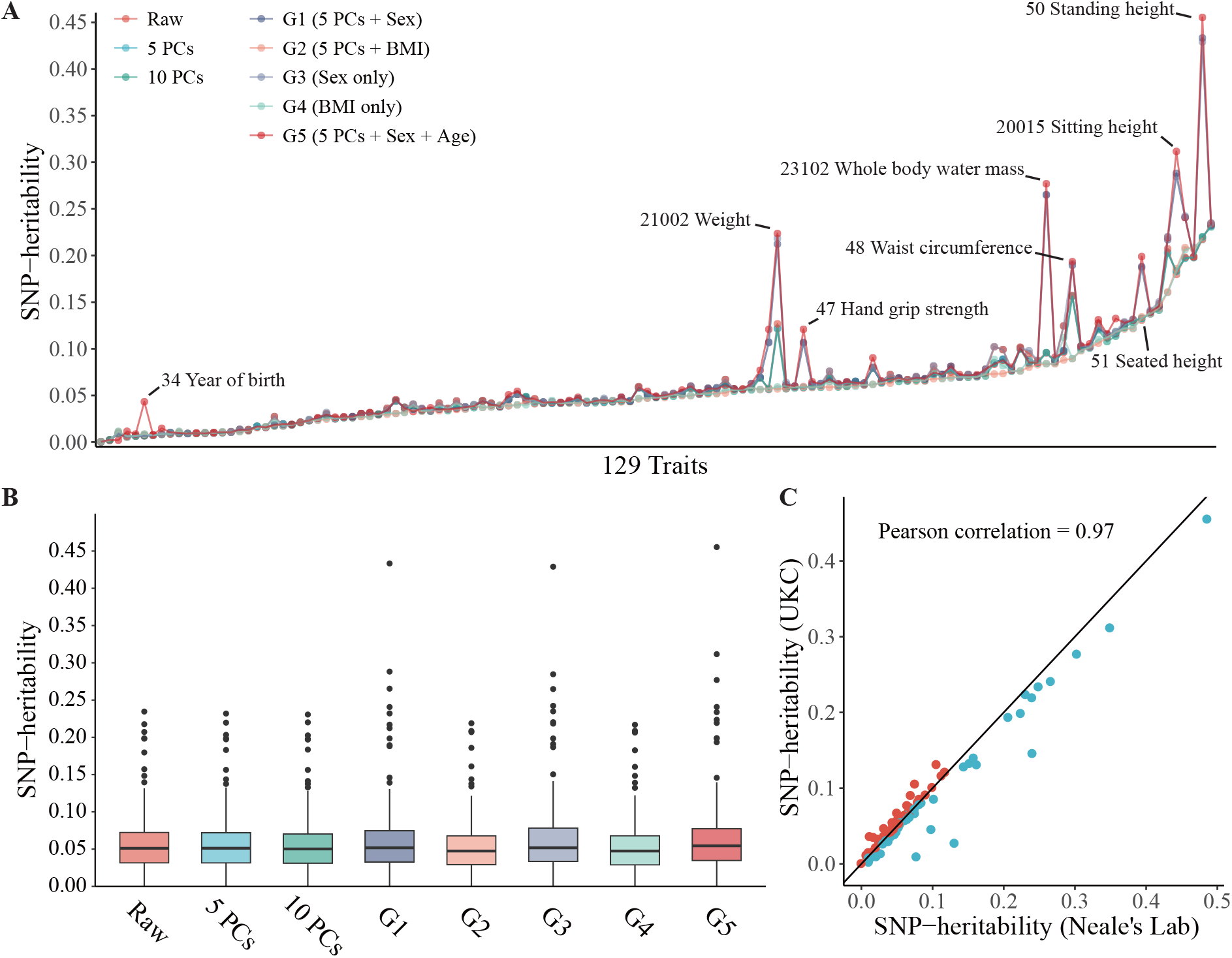
Heritability estimated under 8 sets of covariates corrected. **A**) the SNP-heritability estimated with LD score regression. Eight groups of GWAS summary statistics are generated in UKC. The traits that have different SNP-heritability under different models are annotated. **B)** Average SNP-heritability for 129 traits. **C**) SNP-heritability comparison for 112 traits. Their 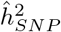 were estimated using UKC, adjusted by 5 PCs, sex and age, and using summary statistics from Neale’s Lab.

Furthermore, we also compared the estimated 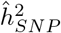 using the UKC summary statistics after adjustment scheme for 5 PCs, Sex, and Age, with 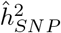 directly downloaded from Neale’s Lab, which was adjusted by sex and the top 10 PCs (UKB heritability, https://nealelab.github.io/UKBBldsc/index.html). Using LDSC, the 112 matched traits had their 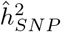 consistently estimated, a Pearson correlation of 0.97 (**Fig. 4 C**). Note that these 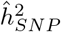 results were all on the observed scale. However, for an ascertainment trait, such as a trait of the case-control design, the prevalence and the relationship between cases and controls should be provided to transform the SNP-heritability from the observed scale to the liability scale [23].

#### 2.3.3 Application 3: Polygenic Score

Polygenic score (PGS), a weighted sum of the number of alleles, measures the risk of the disease based on genetic information [24, 25]. As PGS relies on genetic effects estimated from a GWAS model, the adjustment scheme affects the performance of PGS. We demonstrated how the choice of either UKC-PC or UKB-PC would lead to different results. From the 296,216 unrelated UKB individuals, we randomly selected 10,000 individuals as the test dataset, and the remaining 286,216 individuals as the training dataset. The variants with MAF *<* 0.001, imputation quality score *<* 0.8 or VIF *>* 10 were excluded from the training dataset, and for the test dataset variants with MAF *<* 0.01, missing rate *>* 0.05 or Hardy–Weinberg equilibrium test *p*-value *<* 1e-8, and individuals who had their missing call rate higher than 0.05 were removed. Variants with palindromic alleles between the training and the test datasets were removed. The training model included both Sex and Age as covariates, and the population structure scheme was either corrected by the top 10 UKB-PCs (denoted by **M1**) or the top 10 UKC-PCs (denoted by **M2**). Given the estimated effect 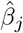 for each SNP *X*_*j*_, the phenotype was predicted by 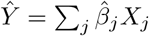 as implemented by “–score” in PLINK [9]. The prediction accuracy was measured by Pearson’s correlation between true phenotype *Y* and 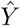 (polygenic genetic score correlation, denoted by *R*) across all test samples, and no further covariates were adjusted for *R*.

The prediction accuracy *R* was evaluated under different sets of *β* after applying *p*-value thresholds, totaling 15 categories ranging from 1e-7 (significant variants) to 1 (all common variants). For each of the 126 traits, we picked the maximum *R* among the 15 categories for **M1** or **M2** adjustments, respectively. The mean *R* were 0.0942 in **M1** and 0.0914 in **M2**, showing no statistical difference (**Fig. 5 A, Supplementary Data V**). However, the PGS results exhibited variation across phenotypic categories. For the phenotypes classified into “Lifestyle and environment”, “Health outcome” and “Mental health”, *R* were stable under different PC adjustments (**Fig. 5 B**). In categories “Physical measurements”, “Family history”, and “Early life factors”, **M1** and **M2** schemes resulted in different *R*. For example, the *R* for Weight was 0.1549 under **M2** but 0.2299 under **M1**. In terms of ‘Family history”, Number of full siblings had a higher *R* under the **M2** than those under **M1** (0.1464 v.s. 0.0659 for Number of full brothers, 0.1111 v.s. 0.0513 for Number of full sisters).

**Figure 5.**
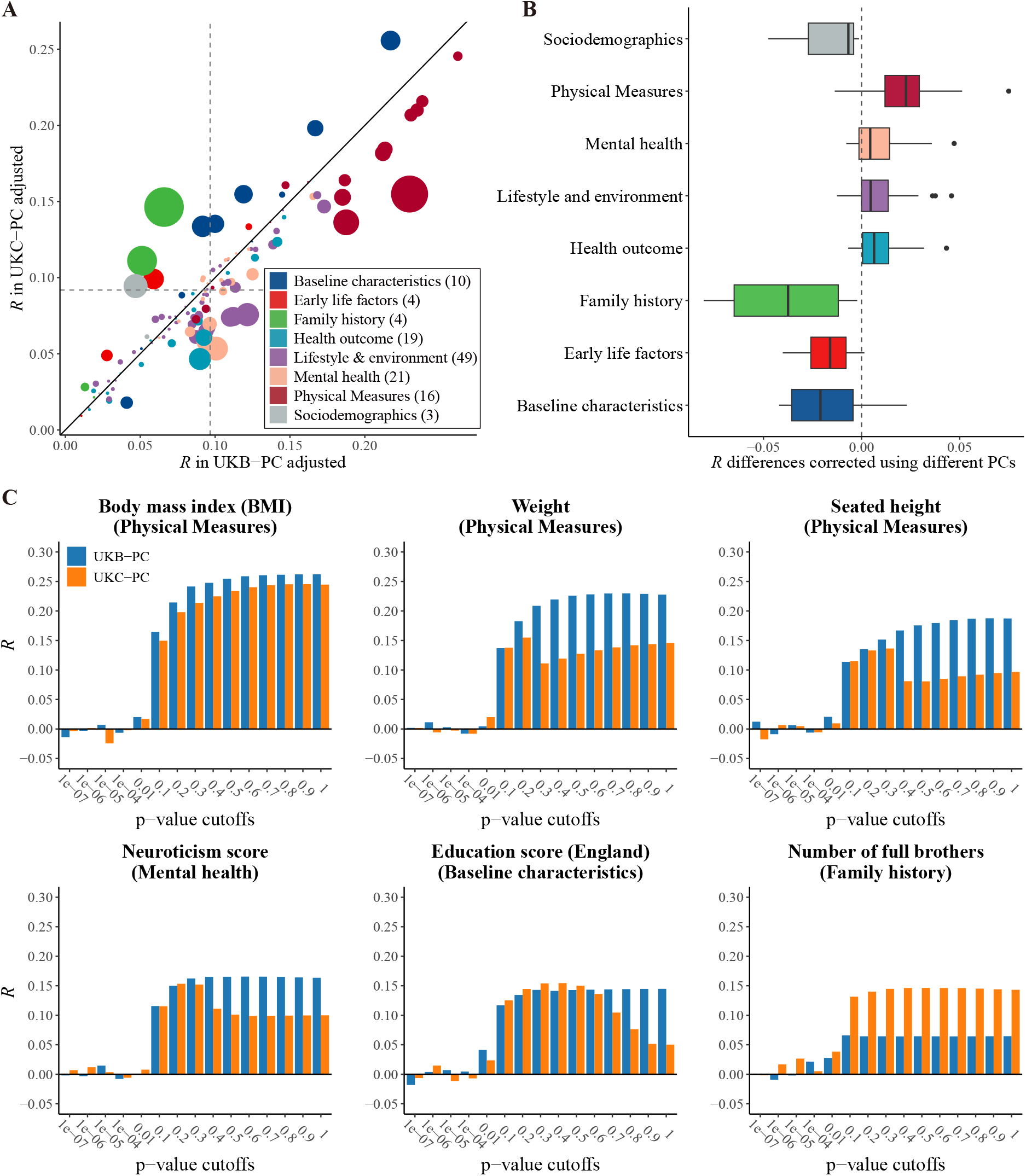
UKC conducts PGS analysis of 126 phenotypes under UKB-PC or UKC-PC adjustments. **A**) Polygenic genetic score correlation (*R*) of all phenotypes corrected by different PCs. The 126 UKB phenotypes were classified into eight categories based on their descriptions. Vertical and horizontal dotted lines for the mean of the 126 *R*. The size of each point is proportional to the difference between two *R* **B**) The distributions of *R* differences (M1-*R* minus M2-*R*) under different categories. **C**) Variation of *R* for the representative traits using variants under different p-value thresholds.

Furthermore, *R* displayed varying trends along the *p*-value thresholds across different phenotypes (**Fig. 5 C**). The *R* of BMI exhibited a consistent increase with larger *p*-value thresholds under both adjustment schemes. On the contrary, Weight, which was highly correlated with BMI, displayed an increasing *R* trend under **M1** but reached its maximum *R* near the *p*-value cutoff at 0.2 under **M2**; a similar trend was observed for Seated height (UKB field ID: 51). For Neuroticism score, its maximum *R* under both adjustment schemes were found near *p*-value thresholds of 0.3. For the Education score (UKB field ID: 26414), its maximum *R* was achieved at *p*-value threshold of 0.4 under the **M1**. Number of full brothers (UKB field ID: 1873) showed a much higher *R* under **M2**.

In this demonstration, the local population structures and cryptic relatedness remained elusive and might influence the performance of PGS. Other factors could also be further investigated using the UKC platform.

#### 2.3.4 Application 4: Mendelian Randomization

Mendelian randomization (MR) is a method used to infer causal effects between exposures and outcomes using genetic variants as instrumental variables (IV) [26]. Two-sample Mendelian randomization is a MR method that utilize estimates of genetic association of outcomes and exposure derived from different samples [27]. In the absence of original data, researchers must rely on existing GWAS summary results that have been adjusted for certain covariates, potentially introducing bias into MR analyzes [28].

To investigate how the adjustment of covariates in GWAS summary statistics could perturb MR results, we used UKC to perform an extensive MR analysis. This involved adjusting for various combinations of covariates to gain a comprehensive understanding of their effects.

We performed covariate-adjusted two-sample MR to investigate the causal relationship between Waist circumference (UKB field ID: 48, WC) and rheumatoid arthritis (RA). We obtained the RA summary statistics from a previous meta-GWAS that included 18 cohorts, consisting of 14,361 RA cases and 43,923 controls of European ancestry [29]. WC summary statistics are generated with UKC adjusting for various combinations of covariates. SNPs with *p*-values *<* 5 *×* 10^*−*8^ underwent linkage disequilibrium clumping(*r*^2^ *<* 0.01 within the distance of clumping 250 kb) were used as IVs in the MR analysis. The inverse-variance weighted (IVW) method as the primary method was used to obtain the estimated effect size, supplemented by other three methods (weighted median estimation, simple median estimation, and MR-Egger regression). We provided an example where MR estimates differed substantially when WC summary statistics were adjusted for different sets of covariates (**Fig. 6, Tab. 2**). In **Fig. 6 A**, the associations between genetic variants and WC were adjusted for BMI and Alcohol intake frequency (UKB field ID: 1558), while in **Fig. 6 B**, the adjustments included Weight, Body fat percentage (UKB field ID: 23099), Smoking status (UKB field ID: 20116), and 10 PCs. Notably, the results revealed a reversal in the direction of estimated effects using IVW and simple median when covariates vary-a phenomenon that had received limited scrutiny but was accessible for thorough investigation through tools like UKC.

**Figure 6.**
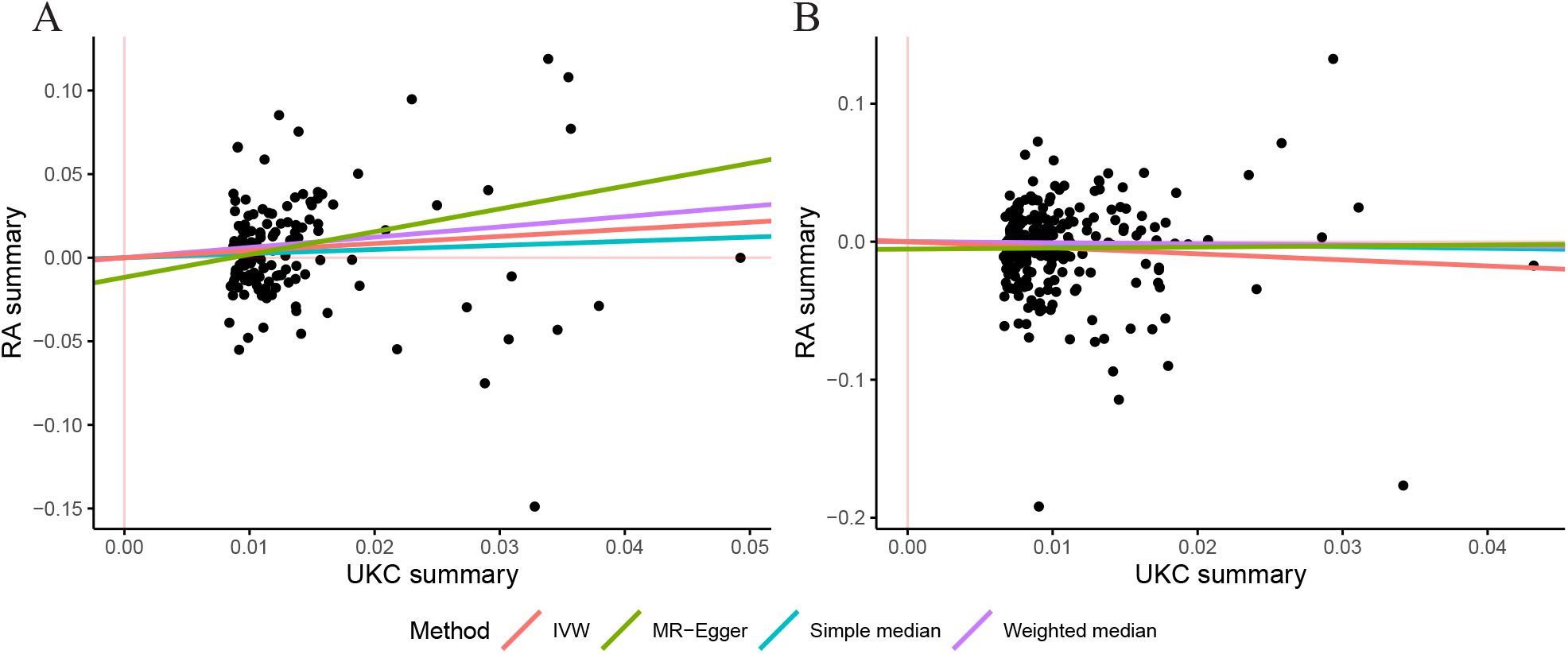
Casual effects of Waist circumference (WC) on rheumatoid arthritis (RA) for different covariates-adjusted two-sample MR studies. **A**) Results adjusted for BMI and Alcohol intake frequency. **B**) Results adjusted for Weight, Body fat percentage, Smoking status and 10 PCs. The x-axis plots the *β* estimates of each SNP on WC. The y-axis plots the *β* estimates of each SNP on RA. The lines in different colors indicate the causal effect estimates by inverse variance weighted, MR-Egger regression, simple median and weighted median methods.

**Table 2:** Summary of casual effects of waist circumference (WC) on rheumatoid arthritis (RA) with different covariates adjusted.

| Covariates | Method | Num. of QTLs | $\beta$ | $\sigma_\beta$ | $p$-values |
| --- | --- | --- | --- | --- | --- |
| BMI &<br>Alcohol intake frequency | Inverse-variance weighted (random) | 138 | 0.423654 | 0.179907 | 0.01853 |
|  | MR-Egger | 138 | 1.363478 | 0.673006 | 0.04277 |
|  | Weighted median | 138 | 0.614955 | 0.240209 | 0.010465 |
|  | Simple median | 138 | 0.245338 | 0.238348 | 0.303326 |
| Weight &<br>Body fat percentage &<br>Smoking status &<br>10 PCs | Inverse-variance weighted (random) | 247 | -0.43919 | 0.170062 | 0.009807 |
|  | MR-Egger | 247 | 0.075531 | 0.570881 | 0.894743 |
|  | Weighted median | 247 | -0.07439 | 0.225956 | 0.74201 |
|  | Simple median | 247 | -0.12095 | 0.226027 | 0.592575 |

¡b¿Casual effects of Waist circumference (WC) on rheumatoid arthritis (RA) for different covariates-adjusted two-sample MR studies.¡/b¿ ¡b¿A¡/b¿) Results adjusted for BMI and Alcohol intake frequency. ¡b¿B¡/b¿) Results adjusted for Weight, Body fat percentage, Smoking status and 10 PCs. The x-axis plots the ¡i¿¡/i¿ estimates of each SNP on WC. The y-axis plots the ¡i¿¡/i¿ estimates of each SNP on RA. The lines in different colors indicate the causal effect estimates by inverse variance weighted, MR-Egger regression, simple median and weighted median methods.

As a proof-of-principle study, we only demonstrate the basic utility of the four applications, and there are other methods to improve their performance [4].

## 3 Availability and Portability

### 3.1 Availability of UK BioCoin

Both the UKC NSS and the UKC computational engine are integrated into a Docker image (20 GB), which can be downloaded from the GitHub repository (https://github.com/Ttttt47/UKBioCoin). As the UKC Docker image has been deployed onto Docker image servers worldwide, it can be successfully downloaded in about an hour as tested in various regions, including Melbourne (Victoria, Australia), Nashville (Tennessee, US), Tokyo (Japan), and Stockholm (Sweden); in mainland China, it takes about 20 minutes to download the UKC Docker image. It should be noted that NSS has been sealed into the UKC image, and the substantial computational cost for NSS (about 2 days for UKB) should not be concerned.

### 3.2 Portablility for Other Biobanks

UKC is not only available as an encapsulated package but is portable to other biobanks, and it is straitforward to build a UKC-like platform. For example, we have successfully applied the entire UKC framework in the Westlake Biobank cohort (WBBC) [10], and have brought out Westlake BioCoin (**WBC**). In this test, WBBC used 5,440 chipped GWAS samples and 14,242,187 QCed SNPs (locus genotyping rate *>* 0.05, HWE *>* 0.00001, MAF *>* 0.001), and it took approximately 42 minutes to convert its original individual-level data (5.06 GB) into the corresponding NSS (1.43 GB). As a validation, WBBC performed individual-level GWAS for height with the inclusion of the top 5 PCs, age, and sex as covariates, and **WBC** yielded, as expected, nearly identical results for the matched *β* and *p*-values. Obviously, the demonstrated four UKC applications, as well as other utilities, can be equivalently conducted for **WBC**. We provide scripts for the conversion of other datasets to establish their own BioCoin like UKC.

## 4 Discussion

Privacy concerns about individual-level data have limited the data availability, precluding the reproducibility of genetic studies and collaboration between biobanks. Public released summary statistics promote data-sharing but lack of flexibility to explore trait-specific covariates, thus narrowing the scope of downstream studies. To address these challenges, we propose a novel framework that facilitates flexible summary statistics data-sharing. Given its pivotal role in providing ingredients for other studies, we select UKB as a working instance and developed UKC, a summary statistics generator integrating UKB and the summary statistics regression technique into a single device. We only cover UKB GWAS analysis, but it can profoundly determine the performance of the estimation of heritability, PGS, and Mendelian randomization, which are highly subject to UKB output.

In order to make UKC highly consistent to UKB GWAS analyses, we require the summary statistics to be generated in the form of naive summary statistics, which are synthesized to carry out nearly exact linear model analysis as individual-level UKB data. As demonstrated, when there is no, low, or even substantially high missing data, UKC continues to deliver high-quality results. Additionally, the quality control metric VIF, which is calculated for each testing SNP, further eliminates the possible bias. After compressing 289 GB UKB source data into 20 GB NSS, UKC is sealed into a portable Docker image, which can be downloaded to a local site in one hour, as tested worldwide. As the computational kernel of UKC works on summary statistics regression, which is further optimized in algorithm and C++ programming, its computational speed is boosted approximately 70 times while requiring little RAM. Therefore, the working environment of UKC can be an average personal laptop.

For UKB GWAS, principal components are most commonly employed covariates. As the correlation matrix of PCs is diagonal, using decomposed inversion of a matrix enables us to derive analytical results for SNP effects and their sampling variance under various possible combinations of PCs. As observed for height, local selection, as captured by EigenGWAS, can lead to high VIF and eventually very obscure GWAS signals. There is no clear clue which set of PCs are suitable for precise mapping of a QTL, but our UKC provides such a device for in-depth evaluation of the stability of GWAS signals, in particular if follow-up experiments are planned to rely on those results. Various adjustments, such as inclusion of sex and age, can be made and their influence has been demonstrated in the application **I**-**IV**.

As a proof-of-principle study, we only include phenotypes commonly employed in UKB studies, and it is possible to include even more phenotypes. For phenotypes of interest but bearing high missing rates, phenotype imputation can be used to improve data quality [30]. The inquiry of GWAS summary statistics can be other emerging biobanks than UKB. The presented framework can be seamlessly applied to Westlake biobank [10], and possibly for other cohorts such as STROMICS [31], ChinaMap [32], All of US cohort [33], and even proteomic data [34]. As enclosed in UKC are summary statistics, it offers a novel route for data-sharing, without hampering data security but harnessing reproducibility and collaboration.

## 5 Materials & Methods

### 5.1 UK Biobank Overview

The UK Biobank (UKB) is a comprehensive database that contains genetic and health information from more than 500,000 participants in the United Kingdom [8]. As a proof-of-principle study, we focus only on the 292,216 unrelated white British for 129 phenotypes, 60 continuous traits, such as height and BMI, and 69 categorical traits, such as sex. Genomic data of about 805,000 markers are collected on all individuals in the cohort, genotype data are then phased and imputed using computationally efficient methods combined with the Haplotype Reference Consortium (HRC) and UK10K haplotype resource. The imputation protocol has increased the number of variants by more than 50 times, to 96 million variants. The genotype data is first imputed and filtered using a minor allele frequency (MAF) cutoff of 0.001 and palindromic SNPs (A/T, G/C biallelic loci), resulting in retention of 10,531,641 SNPs (**Fig. 1 A**, denoted as 10M SNPs), and 488,007 overlap with the chipped SNPs. The phenotype correlation is shown in **Fig. 1 C**, and the average missing rate is 4.1%. It should be noted that the UKB phenotype data may consist of multiple samplings and array data containing multiple data items. To minimize potential biases, we only use the first sampling and, where applicable, the first element of the array for each phenotype. The principal components are generated using 1 million SNPs, which are randomly sampled from the 10M SNPs (UKC-PC); in contrast, the principal components directly downloaded from UKB (UKB-PC) are also included. Otherwise specified, UKC-PC is included for analysis by default.

### 5.2 Westlake Biobank Overview

The Westlake BioBank for Chinese (WBBC) project is a population-based prospective study that recruited a total of *∼*35,000 participants, comprising *∼*28,000 late adolescents with a mean age of 19 and *∼* 7,000 adults older than 65 years, covering 31 provincial administrative regions in China [10, 35, 36]. In this study, 5,492 participants with health (e.g., sex, age, and height) information and SNP array data were included. Specifically, these participants were first genotyped by the high-density Infinium Asian Screening Array. Genotype data were then imputed using the South and East Asian Reference Database (SEAD) reference panel [10]. After phenotype and genotype quality control (–geno 0.05; –hwe 0.00001; –maf 0.001; –mind 0.05), a total of 14,242,187 SNPs and 5,492 participants were retained in the follow-up analysis.

### 5.3 Genome-wide Association Studies

A genome-wide association study (GWAS) executes a regression between the genetic variant *X* and a continuous phenotype *Y* using a linear regression model:

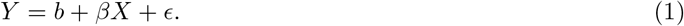

Here, *β* represents the regression coefficient of *X, b* represents the intercept, and *ϵ* constitutes noise following a normal distribution. When *Y* is discrete, a generalized linear model is used to estimate the genetic effect of *X* on *Y*. Assuming 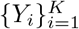 are phenotypes (covariates) measured in a population such as sex and BMI, and 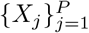 are the numbers of copies of a reference allele with *X*_*j*_ ∈ {0, 1, 2}, 1 *≤ j ≤ P*. Without loss of generality, *X*_*j*_’s are centered to have a mean of zero, while *Y*_*i*_’s are normalized to have a mean of zero and a variance of 1. Generally, the effects of covariates on the phenotype are adjusted to reveal conditional genetic effects, that is, the following model is used to evaluate the genetic effect,

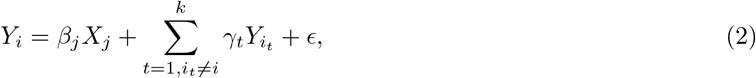

where 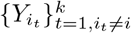 is the set of trait-specific covariates one wants to adjust, and *γ*_*t*_ is the effect of covariate 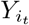 on phenotype *Y*_*i*_.

Furthermore, the population structure is commonly adjusted by including principal components as covariates [19, 20]. Thus, we finally estimate the genetic effect of the SNP using the following model:

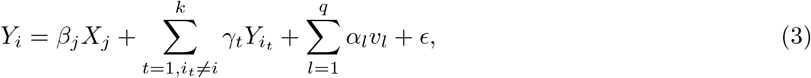

where 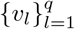 are the top principal components of genetic structure and *α*_*l*_ denotes the regression coefficients of *v*_*l*_. The Ordinary Least Squares (OLS) estimator of the regression coefficient 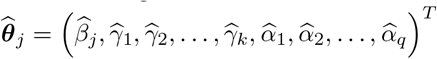 and its estimated variance are given by

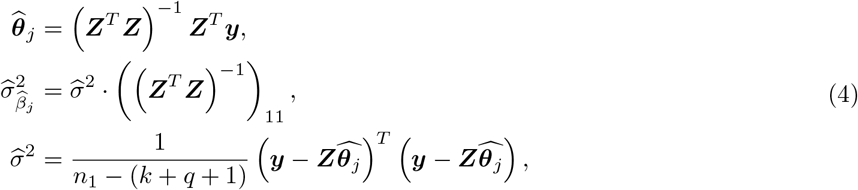

where ***Z*** constitutes an *n*_1_ *×* (*k* + *q* + 1) matrix containing genotype and covariate data of *n*_1_ complete samples, with the *s*^th^ row representing the information of the *s*^th^ sample: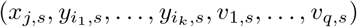, and 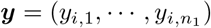 is the observation of phenotype *Y*_*i*_.

### 5.4 UK BioCoin Algorithm

The estimator in **Eq.4** is widely used in GWAS. However, it is not applicable when ***Z*** and ***y*** are not available. We observe that the OLS estimator in **Eq.4** relies on the matrix products ***Z***^*T*^ ***Z, Z***^*T*^ ***y***, and ***y***^*T*^ ***y***, rather than the original data ***Z*** and ***y***. This fact motivates us to use summary statistics regression to get 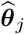 based on summary statistics ***Z***^*T*^ ***Z, Z***^*T*^ ***y***, and ***y***^*T*^ ***y***. Specifically, denote

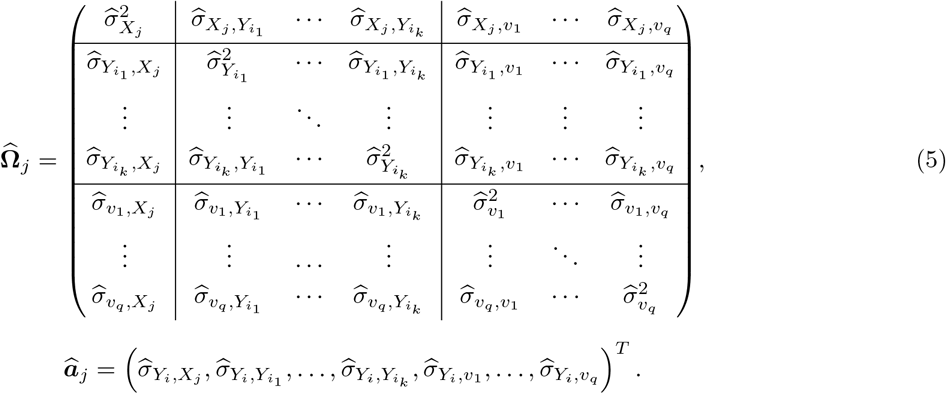

We have

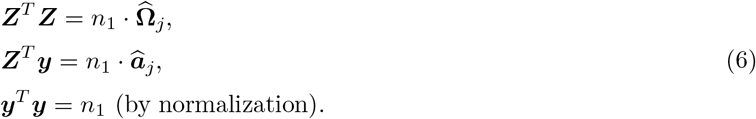

Herein, 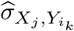 denotes the estimated covariance between *X*_*j*_ and 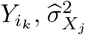 represents the estimated variance of *X*_*j*_, and 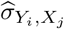 and 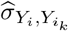 are analogously understood.

Substituting the estimators described in **Eq.6** into **Eq.4** and following a series of elementary calculations, we arrive at the estimators:

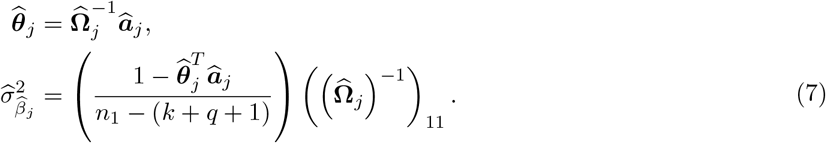

Although these estimators appear to be concise in form, it is important to recognize that in the presence of missing SNP and phenotype data, it is not feasible to obtain 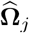 and 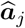. This is due to the fact that the set of complete samples depends on the specific model established, which is unknown beforehand.

Let 𝒮= {*S*_*i*_ = (*x*_1,*i*_, …, *x*_*P,i*_, *y*_1,*i*_, …, *y*_*K,i*_) : *i* = 1, …, *n}* be the entire set of observations, where some of them may contain missing value. At first sight, we can estimate 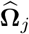 and 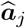 based on 𝒮_0_, where

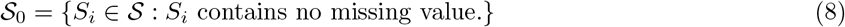

is obtained by discarding all samples that have missing values. However, after quality control we find that none of the samples have complete observations in all SNPs and phenotypes, i.e. 𝒮_0_= *∅*, which makes this approach impracticable. Looking inside the problem, we note that the elements of 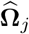 and 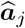 depend only on pairs of variables rather than all of them. This fact suggests to estimate the element 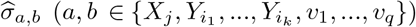 of 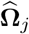 based on samples with complete observations on (*a, b*), which gives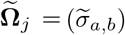. Vector 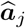 can be estimated in a similar way, denoted by 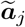.

It should be recognized that the (complete) samples for estimating 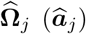 constitute only a subset of samples used in calculating any entries in 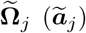 and the distribution of SNPs or phenotypes may differ between these two sets. Therefore, we need to control the missing rates of the covariates included in the analysis to reduce the effects of unbalanced missing pattern and thus the risk of biased estimation of 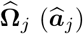.

Subsequently, we approximate the complete sample size *n*_1_ with 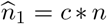, where *c* is a constant that provides a rough approximation of the overall non-missing rate, and *n* is the known total sample size. In practice, one can choose *c* as the product of non-missing rates of phenotypes/SNPs selected in the model, assuming that the absence of these variables is independent of each other, or simply set *c* = 1 when the data is nearly complete. In our implementation, we adopt the former method, that is, 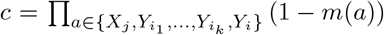, where *m*(*a*) is the missing rate of variable *a*.

Substituting 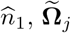 and 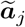 into the **Eq.7** yields the final estimators:

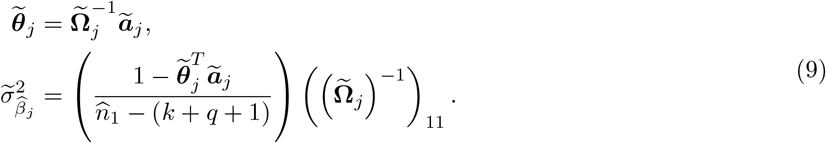

We now examine all conceivable models that could emerge in **Eq.3**, where 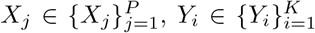 and 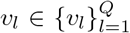. Following the identical estimation procedure delineated above, we discern that the entries of 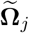 and 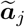 for estimating each model are, in fact, reusable. Indeed, for any potential model in the form of **Eq.3**, UK BioCoin relies exclusively on a set of these entries. To simplify the notation. we logically reorganize it into the subsequent three components:

I. 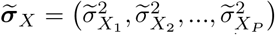, a vector of length *P* that represents the estimated variances of the *P* SNPs.
II. 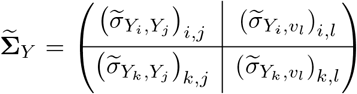, a (*K* + *Q*) *×* (*K* + *Q*) matrix represents the correlation coefficients between the *K* phenotype and *Q* principal components.
III. 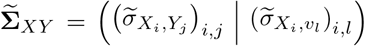, a *P ×* (*K* + *Q*) matrix represents the covariance between the *P* SNPs and (*K* + *Q*) phenotype and principal components.

In addition, if one wants to estimate the overall non-missing rate *c*, a vector describing missing rate of all *P* SNPs and *K* phenotypes is required:

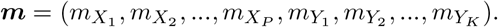

We refer to these statistics 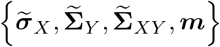 as UKB Naïve Summary Statistics (NSS) in the sense that the UKC estimation are solely based on these statistics. The comprehensive process of UKC is delineated in **Fig. 1**.

### 5.5 Generating Naive Summary Statistics

We will now outline the process of generating the NSS for a given dataset, which serves as a prerequisite for the UKC platform. It is important to note that this procedure needs to be executed only once for a specific dataset.

To generate NSS, we first perform quality control on the raw data and then generate principal components (PCs) from the genotype data to approximate the population structure. These PCs, combined with phenotypes, are subsequently scaled to have unit variance and a zero mean. It should be emphasized that while we also assumed in the previous section that every *X*_*j*_ has a mean of zero, centering the genetic data is not required for generating NSS because the NSS is invariant to mean shifting.

The second step is to calculate the variance for all SNPs presented in the genotype data. To achieve this, for each SNP, we count the frequencies of the three genotypes: *p*_*AA*_, *p*_*Aa*_, and *p*_*aa*_. The variance of a SNP is calculated as 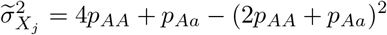. Subsequently, we compute 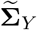 element-wise. The estimate 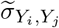 is given by 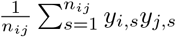, where *y*_*i,s*_ denotes the *i*^th^ phenotype value of the *s*^th^ sample and *n*_*ij*_ denotes the number of complete pairs of observations. The estimates 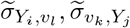 and 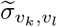 are calculated analogously.

Lastly, we need to compute 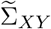. Although this can be achieved by directly estimating the covariance between *X*_*j*_ and *Y*_*i*_ in the same way as the estimation procedure for 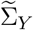, the computational burden for datasets with tens of millions of SNPs, such as UKB, is considerable. To improve computational efficiency, we choose an indirect method to calculate 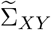. In particular, we first need to perform a single-variable linear regression on every phenotype and principal component. Specifically, we use the following model in PLINK [9]:

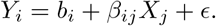

Here, *Y*_*i*_ ∈ {*v*_1_, *v*_2_, …, *v*_*Q*_, *Y*_1_, …, *Y*_*k*_}, *X*_*j*_ ∈ {*X*_1_, *X*_2_, …, *X*_*p*_}, *b*_*j*_ is the intercept and *ϵ* is the noise. We now obtain the estimated regression coefficient 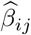, from which 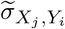 is calculated by

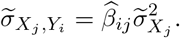

By synthesizing these elements and the missing rates profile, we construct the naïve summary statistics: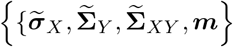.

### 5.6 Estimation of VIF

The variance inflation factor (VIF) for testing the *j*^th^ SNP is defined as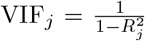, where 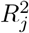 stands for the proportion of variance in *X*_*j*_ that could be explained by the other covariates. VIF reflects the degree of variance inflation of the regression coefficient estimator 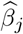 in the sense that it is a factor in the estimated variance 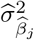 [37]:

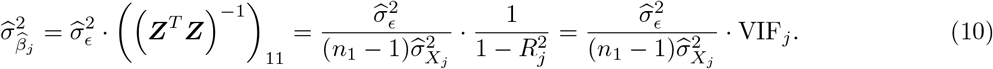

This suggests VIF as a measure of sensitivity of estimate 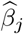 to the variation in the data. SNPs with high VIF are often removed from the results in the sense that they have rather unstable estimates.

In practice, we substitute 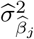 by 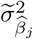 in **Eq.9** and the VIF of the *j*^th^ SNP is given by

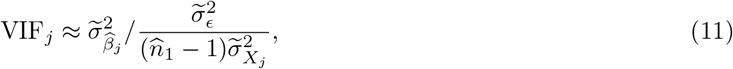

where 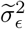 is the mean squared error and the estimator of the variance of the error term *ϵ*:

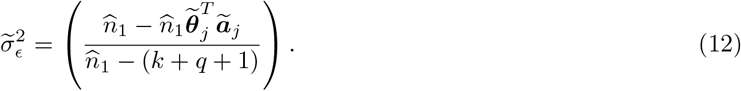

The principal components are widely used covariates in GWAS. When all covariates are 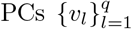, the relationship between VIF and regression is more straightforward. In such a case, since the PCs are independent from each other, 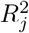 is essentially the sum of squared correlations between *X*_*j*_ and the PCs,

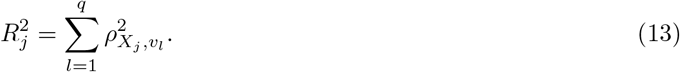

Such correlations between genetic variants and PCs can be revealed by EigenGWAS analysis. EigenGWAS is a flexible genomic scan method to find loci under natural selection[14, 38], which is done in the same manner as GWAS, replacing the phenotype *Y* with PC *v*_*q*_ as the response variable,

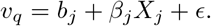

A significant EigenGWAS signal corresponds to a significant correlation 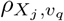 between the SNP and the specific PC, which eventually leads to inflated 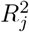 and VIF when adding this PC as covariates in a GWAS. It is worth noting that all PCs form an orthonormal basis of span (*X*_1_, *X*_2_, …, *X*_*P*_), allowing *X*_*j*_ to be represented as a linear combination of *v*_*l*_’s. Consequently, we view 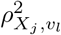 as the inner product of *X*_*j*_ and *v*_*l*_, implying that as more PCs are added as covariates, 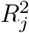 tends toward 1 and VIF tends to +*∞*. This leads to severe multicollinearity and obscure results. Therefore, the selection of the number of PCs is a trade-off between avoiding multicollinearity and correcting for population structure.

When all covariates are PCs, one can also derive the OLS estimator for the regression coefficient for the *j*^th^ SNP *β*_*j*_ as well as the *t* -statistic *t*_*j*_ as

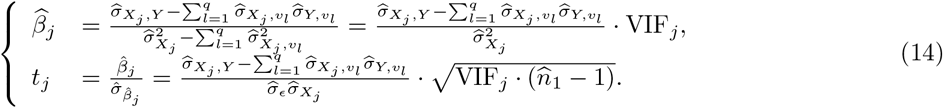

Again, these equations suggest VIF as a measure of stability in the sense that small errors in estimation of 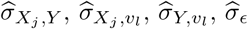 and 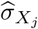 will be amplified by large VIF:

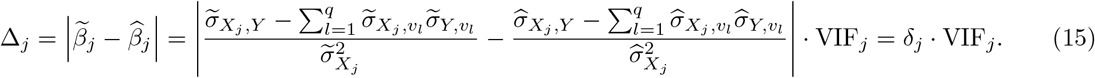

Due to the approximations used in UKC, more errors are introduced compared to individual-level data-based methods. We suggest using a stringent VIF threshold to exclude estimates that not only have high variation but also have a high risk of amplifying the errors introduced by UKC approximation.

## Supporting information

ExtendData

## 6 Data Availability

Westlake Biobank: https://wbbc.westlake.edu.cn

UK Biobank: http://www.ukbiobank.ac.uk/

Neale’s Lab: https://nealelab.github.io/UKBBldsc/index.html

LDSC: https://github.com/bulik/ldsc

PLINK: http://www.cog-genomics.org/plink2/

## 7 Code Availability

UK BioCoin: https://github.com/Ttttt47/UKBioCoin

## 8 Acknowledgements

We thank the participants of UK Biobank (UKB application 41376) and the participants of Westlake Biobank for making this work possible. This work was partially supported by Key R&D Program of Zhejiang Province (2021C03G2013079 to HJ), National Natural Science Foundation of China (31771392 to CGB, 82174208 and 81973663 to YM, and 32061143019 and 82370887 to HFZ), CNTC (110202101032 (JY-09) to HJ and GBC), and GZY-ZJ-KJ-23001 to GBC. Sponsors did not play a role in the design, preparation, and submission of the article. We gratefully acknowledge the support of high-performance computing from the Center for Bioinformatics and Big Data Technology at Zhejiang University and the High-performance Computing Center at Westlake University. We thank Yun Liu and Yuchao Yu for their helpful discussions and assistance in making this research possible. Many thanks to our friends, who live on different continents, for testing UKC.

